# Developmental Regulation of Homeostatic Plasticity in Mouse Primary Visual Cortex

**DOI:** 10.1101/2021.06.11.448148

**Authors:** Wei Wen, Gina G. Turrigiano

## Abstract

Homeostatic plasticity maintains network stability by adjusting excitation, inhibition, or the intrinsic excitability of neurons, but the developmental regulation and coordination of these distinct forms of homeostatic plasticity remains poorly understood. A major contributor to this information gap is the lack of a uniform paradigm for chronically manipulating activity at different developmental stages. To overcome this limitation, we utilized Designer Receptors Exclusively Activated by Designer Drugs (DREADDs) to directly suppress neuronal activity in layer (L) 2/3 of mouse primary visual cortex (V1) at two important developmental timepoints: the classic visual system critical period (CP, P24-29), and adulthood (P45-55). We show that 24 hours of DREADD-mediated activity suppression simultaneously induces excitatory synaptic scaling up and intrinsic homeostatic plasticity in L2/3 pyramidal neurons during the CP, consistent with previous observations using prolonged visual deprivation. Importantly, manipulations known to block these forms of homeostatic plasticity when induced pharmacologically or via visual deprivation also prevented DREADD-induced homeostatic plasticity. We next used the same paradigm to suppress activity in adult animals. Surprisingly, while excitatory synaptic scaling persisted into adulthood, intrinsic homeostatic plasticity was completely absent. Finally, we found that homeostatic changes in quantal inhibitory input onto L2/3 pyramidal neurons were absent during the CP but present in adults. Thus, the same population of neurons can express distinct sets of homeostatic plasticity mechanisms at different development stages. Our findings suggest that homeostatic forms of plasticity can be recruited in a modular manner according to the evolving needs of a developing neural circuit.

**Significance statement:** Developing brain circuits are subject to dramatic changes in inputs that could destabilize activity if left uncompensated. This compensation is achieved through a set of homeostatic plasticity mechanisms that provide slow, negative feedback adjustments to excitability. Given that circuits are subject to very different destabilizing forces during distinct developmental stages, the forms of homeostatic plasticity present in the network must be tuned to these evolving needs. Here we developed a method to induce comparable homeostatic compensation during distinct developmental windows, and found that neurons in the juvenile and mature brain engage strikingly different forms of homeostatic plasticity. Thus, homeostatic mechanisms can be recruited in a modular manner according to the developmental needs of the circuit.

## Introduction

Homeostatic plasticity ensures the proper development of neocortical circuitry by maintaining network stability (Turrigiano and Nelson, 2004). Past works show that neocortical pyramidal neurons utilize three main routes of homeostatic control: modulation of excitation, inhibition, and intrinsic excitability (Turrigiano, 2011). Evidence suggests that these mechanisms are sometimes simultaneously engaged (Lambo and Turrigiano, 2013; Wu et al., 2020), but other times are not (Maffei and Turrigiano, 2008b; Barnes et al., 2015). Therefore, it remains unclear how they are coordinated within the network. Furthermore, our understandings of homeostatic mechanisms come primarily from the malleable juvenile brain, and information about how these mechanisms evolve as neural circuits mature is scarce. A major impediment to answering these questions has been the lack of a uniform activity manipulation method to induce homeostatic plasticity across development. Here we report a reliable chemogenetic approach that allows us to manipulate the same population of neurons *in vivo* at distinct developmental stages, and use this method to characterize the developmental regulation of homeostatic plasticity within L2/3 of mouse V1.

Many forms of homeostatic compensation have been described in a variety of neuronal circuits and cell types (Zhang and Linden, 2003; Marder and Goaillard, 2006; Turrigiano, 2012; Davis, 2013). In neocortical networks, synaptic and intrinsic homeostatic plasticity are widely expressed; the former adjusts synaptic inputs in the appropriate direction to compensate for changes in activity, while the latter modulates intrinsic excitability to alter the input-output relationship of the neuron (Turrigiano, 2011). These mechanisms were first identified in dissociated neuronal cultures, where activity can be easily perturbed but developmental changes are difficult to assess (O’Brien et al., 1998; Turrigiano et al., 1998; Desai et al., 1999; Burrone et al., 2002). Subsequent studies in sensory cortex used an “ex vivo” approach in which sensory deprivation was followed by acute slice recordings (Maffei and Turrigiano, 2008a; Gainey and Feldman, 2017). This allows assessments of homeostatic plasticity in intact networks, but several challenges remain. First, sensory deprivation paradigms generally induce both Hebbian and homeostatic plasticity in parallel, so disentangling one mechanism from another can be challenging. Second, the same paradigm can have fundamentally different effects on the patterns of activity reaching cortex at different developmental stages, making it difficult to directly compare homeostatic compensation across development. Recently, DREADDs have emerged as an alternative for activity manipulation *in vivo* (Roth, 2016). Using this toolset, we can directly suppress activity in a specific cell type at particular developmental stages, and thus assess how homeostatic mechanisms are developmentally regulated.

In this study, we unilaterally expressed inhibitory DREADD hM4Di in L2/3 pyramidal neurons in monocular mouse V1 (V1m), and delivered clozapine dihydrochloride (CNO) to the animal to suppress the activity of DREADD-expressing neurons for 24 hours. We recorded miniature excitatory postsynaptic currents (mEPSCs) from both hM4Di-positive neurons in the injected hemisphere and hM4Di-negative neurons in the control hemisphere. During the classical visual system CP, we found that hM4Di activation scaled up mEPSC amplitude and increased intrinsic excitability. Notably, DREADDs-induced synaptic scaling was blocked by the C-terminal tail (C-tail) of GluA2 AMPA receptor subunit, as is pharmacologically and sensory-deprivation-induced synaptic scaling (Gainey et al., 2009; Lambo and Turrigiano, 2013). Further, this same paradigm failed to induce synaptic scaling and intrinsic homeostatic plasticity in Shank3 knockout (KO) mice, which are known to have deficits in both forms of homeostatic plasticity (Tatavarty et al., 2020). Interestingly, although activity-suppression in adult V1 continued to elicit robust synaptic scaling, intrinsic homeostatic plasticity was absent. Recordings of miniature inhibitory postsynaptic currents (mIPSCs) from adult but not CP neurons revealed a reduction in frequency, suggesting that the mature cortex utilizes inhibitory mechanisms in lieu of intrinsic homeostatic plasticity. Our data show that the same population of neurons can engage different sets of homeostatic mechanisms at distinct developmental stages. We propose that excitatory, inhibitory, and intrinsic homeostatic plasticity mechanisms subserve modular functions, and can be turned on and off to suit distinct developmental needs.

## Materials and Methods

### Key resource table

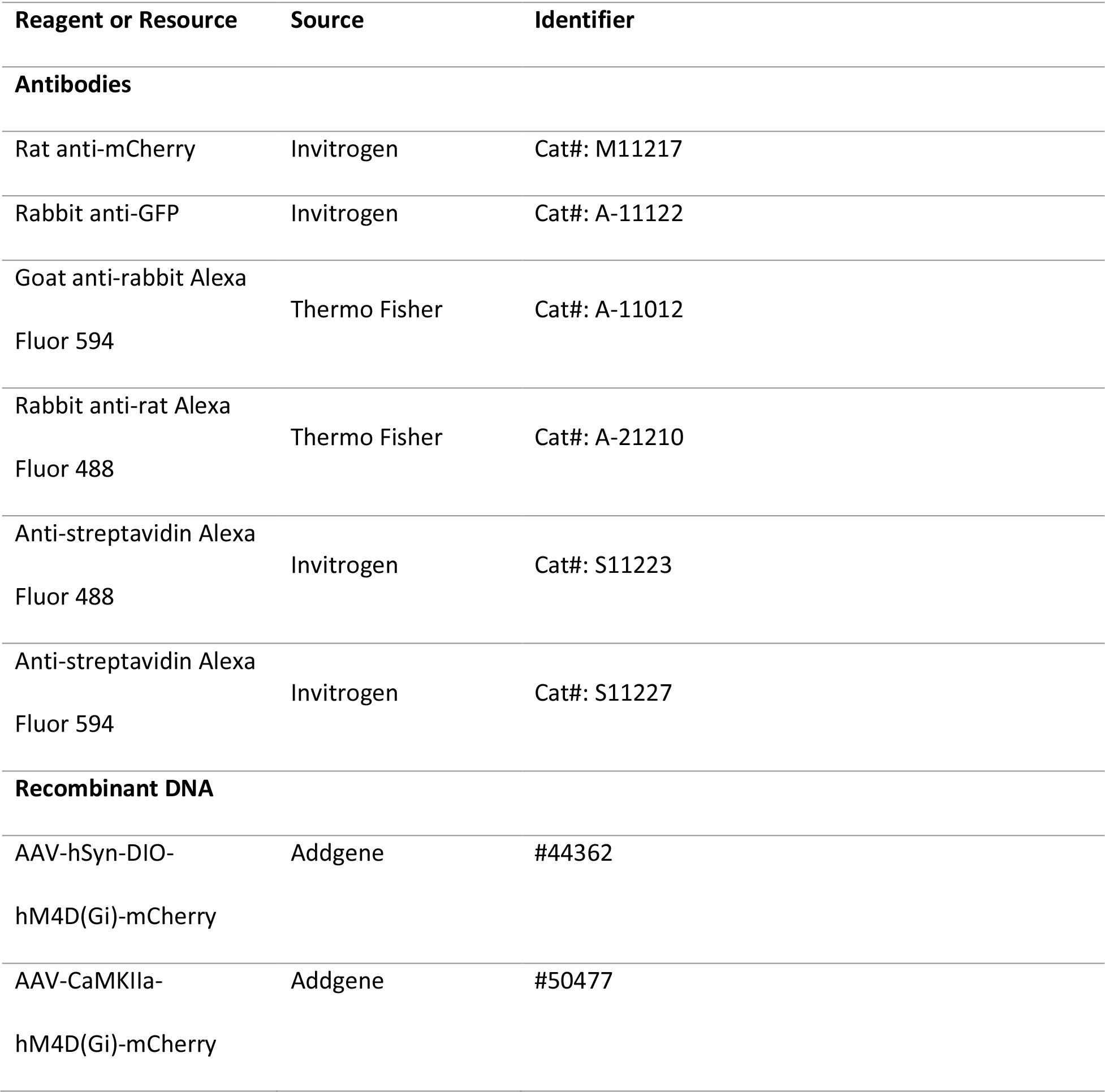
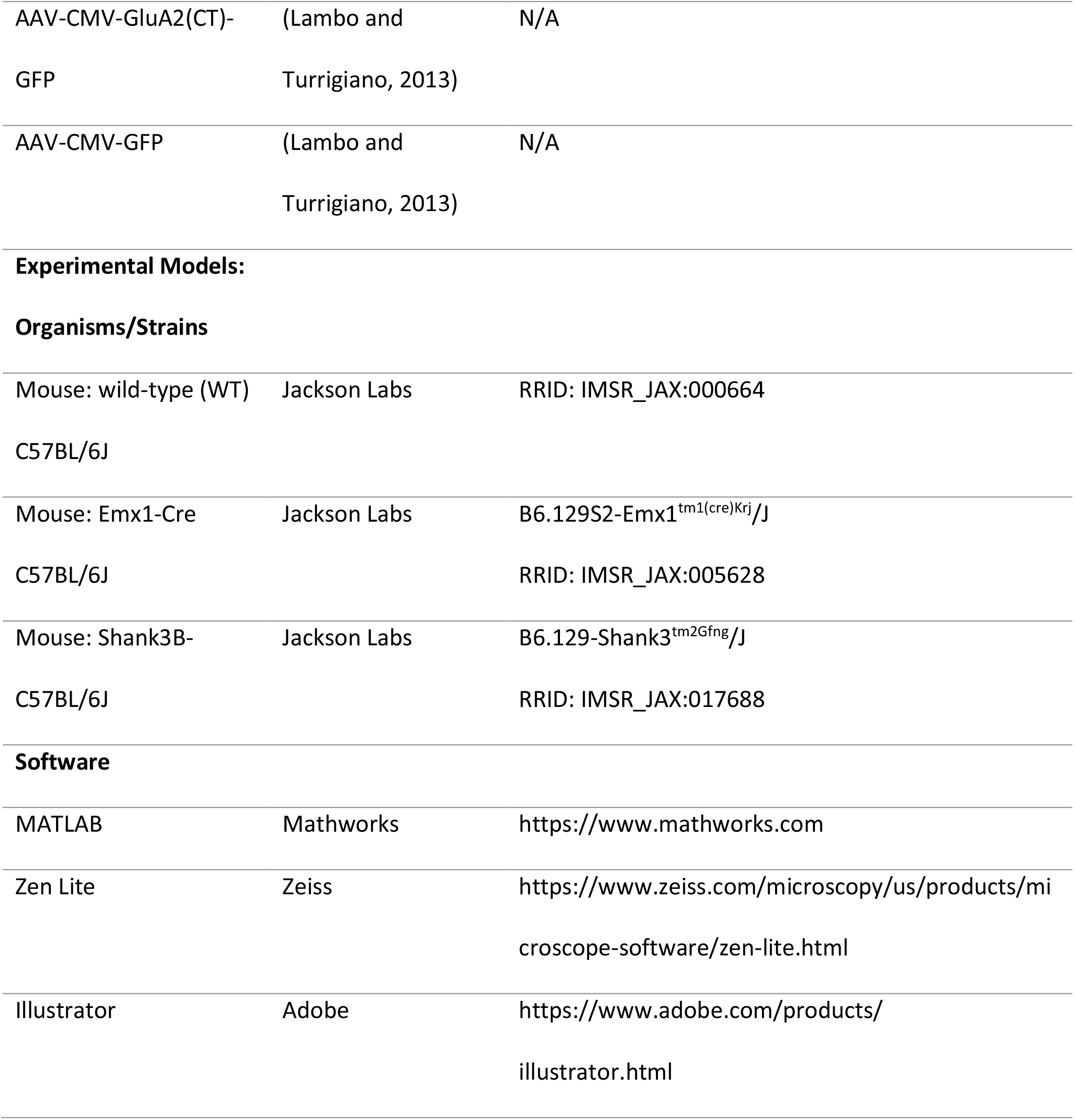

### Animals

Experiments were performed on C57BL/6J mice (both WT and transgenic mouse lines). For all experiments, both males and females were used for slice physiology, either between postnatal day (P) 24 and 29 (for juveniles) or 45 and 55 (for adults). All animals were treated in accordance with Brandeis IBC and IACUC protocols. Specifically, they were housed on a 12:12 light/dark cycle with *ad libitum* access to food and water except when experiments dictated otherwise (see below for details). Pups were weaned between P19 and P21 and housed with at least one littermate except when experiments required single-housing. For most experiments, genetic manipulations (e.g., virus injection) were done unilaterally so that the other hemisphere could serve as a within-animal control. The number of animals and neurons for each experiment are given in the figure legends, and individual data points represent neurons unless indicated otherwise.

### Drug administration

For intraperitoneal injections, clozapine dihydrochloride (CNO, Hello Bio, Princeton NJ) was dissolved in 0.9% sterile saline to reach the desired concentration (1 mg/mL). CNO was then administered every 12 hours, starting 24 hours before slice physiology (two injections in total) at a dose of 5 mg/kg. For drinking water administrations, CNO was dissolved in water to 0.05 mg/mL, and 10 mM saccharine chloride was added to the solution before giving to the animals. All animals that underwent the drinking water paradigm were water-deprived for 16 hours before switching the regular water to the CNO-containing water. They were sacrificed for slice physiology after 24 hours of CNO administration unless indicated otherwise.

### Virus injection

Most virus injections were performed between P14 and P19 on a stereotaxic apparatus under ketamine/xylazine/acepromazine anesthesia. For adult slice physiology experiments, virus injections were performed between P20 and P22. V1m was targeted unilaterally using stereotaxic coordinates (Allen Brain Atlas) that were proportionally adjusted according to the age-dependent bregma-lambda distance difference. Unless noted otherwise, 200-300 nL of virus were delivered into the targeted area via a micropipette. Surgerized animals were allowed to recover in their home cages for a week before slice physiology experiments.

### Ex vivo brain slice preparation

Animals were anesthetized with isoflurane. After toe-pinch check, the animal was decapitated and coronal slices (300 µm) containing V1m from both hemispheres were obtained. Slices were first transferred to an oxygenated chamber filled with choline solution (in mM: 110 Choline-Cl, 25 NaHCO_3_, 11.6 Na-Ascorbate, 7 MgCl_2_, 3.1 Na-Pyruvate, 2.5 KCl, 1.25NaH_2_PO_4_, and 0.5 CaCl_2_, osmolarity adjusted to 310 mOsm with dextrose, pH 7.4) for recovery, and then transferred back to oxygenated standard artificial cerebrospinal fluid (ACSF, in mM: 126 NaCl, 25 NaHCO_3_, 3 KCl, 2 CaCl_2_, 2 MgSO_4_, 1 NaH_2_PO_4_, 0.5 Na-Ascorbate, osmolarity adjusted to 310 mOsm with dextrose, pH 7.4) and incubated for 40 min. Slices were used for electrophysiology 1-5 hours post slicing.

### Electrophysiology

V1m was identified in acute slices using white matter morphology as a guide. Pyramidal neurons were visually targeted and identified by their teardrop shaped somata and the presence of an apical dendrite, followed by post hoc confirmation from biocytin fill reconstruction. DREADD-positive neurons were identified by expression of fluorescent markers. Borosilicate glass pipettes were pulled to achieve pipette resistances between 4 to 6 MΩ and were filled with K^+^ Gluconate-based internal solution (in mM: 100 K-gluconate, 10 KCl, 10 HEPES, 5.37 Biocytin, 0.5 EGTA, 10 Na-Phosphocreatine, 4 Mg-ATP, and 0.3 Na-GTP, osmolarity adjusted to 295 mOsm with sucrose, pH adjusted to 7.4 with KOH) unless otherwise noted. All recordings were performed in slices that were superfused in oxygenated standard ACSF at 34°C. Neurons were visualized on an Olympus BX51WI upright epifluorescence microscope using a 4x air and a 40x water-immersion objective with infrared-DIC optics. Data were acquired at 10 kHz with Multiclamp 700B amplifiers and CV-7B headstages (Molecular Devices, Sunnyvale CA) unless otherwise noted; mEPSC/IPSC data were low-pass filtered at 5 kHz, intrinsic excitability data were not filtered. Data were acquired using the open-source MATLAB-based software WaveSurfer (HHMI Janelia, Ashburn VA), and data analysis was performed using in-house scripts written in MATLAB.

#### *mEPSC recording*s

For spontaneous mEPSC recordings, slices were superfused with standard ASCF containing a drug cocktail of tetrodotoxin (TTX, 0.1 µM), D-2-amino-5-phosphonovalerate (AP-V, 50 µM), and picrotoxin (25 µM) to isolate mEPSCs. L2/3 pyramidal neurons were targeted and held at -70 mV in whole-cell voltage clamp. Each neuron was recorded for 3-5 minutes in a series of 30s segments, and a 500 ms 5mV hyperpolarizing voltage step was administered at the beginning of each segment so that passive properties could be monitored throughout the recording. Neurons were excluded if access resistance was > 20 MΩ, input resistance was < 100 MΩ, membrane potential was > -55 mV, or these properties changed by > 15% during the recording.

#### Intrinsic excitability measurement

For intrinsic excitability measurements, slices were superfused with standard ACSF containing AP-V (50 µM), picrotoxin (25 µM), and 6,7-dinitroquinoxaline-2,3-dione (DNQX, 25 µM) to block synaptic currents. L2/3 pyramidal neurons were held in current clamp with a small DC current injection to maintain the resting membrane potential at -70 mV. Frequency versus current (f-I) curves were obtained by delivering a series of 20 1s long current injections in amplitude increments of 20 pA. Passive properties were monitored in voltage clamp before and after current steps, and neurons that did not meet the criteria listed above for mEPSC recordings were excluded; in addition, neurons were excluded if the deviation of baseline potential from -70 mV was > 5%.

#### mIPSC recordings

For spontaneous mIPSC recordings, slices were superfused with standard ASCF containing TTX (0.1 µM), AP-V (50 µM), and DNQX (25 µM) to isolate mIPSCs. L2/3 pyramidal neurons were held at -70 mV under voltage clamp. The internal solutions for these experiments were adjusted so that the reversal potential for Cl^-^ was 0 mV (in mM: 120 KCl, 10 HEPES, 5.37 Biocytin, 0.5 EGTA, 10 Na-Phosphocreatine, 4 Mg-ATP, and 0.3 Na-GTP, osmolarity adjusted to 300 mOsm with sucrose, pH adjusted to 7.4 with KOH) and mIPSCs were recorded as inward currents. Inclusion criteria were adjusted for these recording conditions: neurons were excluded if access resistance was > 20 MΩ, input resistance was < 90 MΩ, membrane potential was > -60 mV, or passive properties changed during the recording by > 15%.

#### Spontaneous firing

To measure spontaneous firing, slices were washed into active ACSF (in mM: 126 NaCl, 25 NaHCO_3_, 3.5 KCl, 1 CaCl_2_, 0.5 MgCl_2_, 0.5 NaH_2_PO_4_, 0.5 Na-Ascorbate, osmolarity adjusted to 310 mOsm with dextrose, pH 7.4). Whole-cell recordings from DREADD-expressing L2/3 pyramidal neurons were obtained in current clamp. After 5 min of recording, active ACSF containing 500 nM CNO was washed in. Passive properties were monitored before and after wash-in as described above. Neurons were excluded if access resistance was > 20 MΩ, input resistance was < 100 MΩ, membrane potential was > -55 mV, or any property during the recording changed by > 15%.

### Slice immunohistochemistry

Following slice physiology recordings, slices were post-fixed in 4% PFA overnight and transferred to PBS for storage before staining. Free-floating slices were then blocked with 1% BSA/PBS (with 0.1% Triton-X and 0.05% NaN_3_) at room temperature for 2 hours. Slices were incubated with primary antibodies at 4 °C overnight, and then with secondary antibodies at 4 °C overnight. After washing, slices were mounted in Fluoromount G mounting medium (Southern Biotech, AL) and images were obtained using Zeiss LSM880 confocal microscope (Zeiss, Oberkochen Germany).

### Quantification and statistical analysis

Data were analyzed using in-house scripts written in MATLAB. For each experiment, unless otherwise noted, results are reported as mean ± S.E.M. and effect sizes are reported as a percentage of control; sample sizes (both the number of neurons and number of animals), statistical tests used, and p values are given in either the corresponding results section or the figure legends.

#### mEPSC/IPSC recordings

Mean amplitude and frequency were first calculated for each event for 30 s of recording values were then averaged to give the mean value for each neuron. Rise time and decay time constants were calculated from the waveform average traces for each neuron. Rise time is defined as the time for the current to increase from 10% to 90% of the peak amplitude. For mEPSC recordings, decay time constant (τ) is derived from a first-order exponential fit of the decay phase. For mIPSC recordings, the decay phase is fitted to a second-order exponential function, yielding a fast and slow decay τ (decay_fast, decay_slow), respectively. The percentage of the fast component (fast%) is calculated at the peak. To generate the cumulative distribution function (CDF) of mean amplitudes for each condition, 100 events were randomly selected from each neuron and pooled.

#### Intrinsic excitability recordings

Definitions of neuronal properties and firing parameters were primarily adapted from the electrophysiology technical white paper published by Allen Institute (https://celltypes.brain-map.org/). Briefly, mean instantaneous firing rate (IFR) is defined as the mean reciprocal of the first two interspike intervals. Spike adaptation index is defined as the sum of interspike intervals during a 300 pA current step; higher values indicate more adaptation. Latency is defined as the time difference between the current injection onset and the first spike. Spike width is defined as the width at half maximum height for the first action potential evoked at the rheobase. Action potential voltage threshold is defined as the membrane potential when dV/dt reaches 5% of the maximum dV/dt during the depolarizing phase of the first action potential evoked at rheobase; this measurement can produce more consistent estimates of threshold across cells of the same type compared to the one based on an absolute dV/dt value (Jackson et al., 2004). Afterhyperpolarization (AHP) amplitude is calculated for the first action potential evoked at rheobase, defined as the difference between the minimum membrane potential reached after the action potential and the average membrane potential between the current injection onset and the firing threshold; AHP amplitudes from different cells are normalized to the peak amplitude of the same cell.

#### Statistical analysis

All data sets were subjected to a normality test (Shapiro-Wilk test). For normally distributed data, an unpaired two-sample t test was used for pairwise comparison, a one-way ANOVA followed by Tukey post hoc correction was used for multiple comparisons (no. of groups > 2). For other data, a Mann-Whitney U test was used for pairwise comparison, and a Kruskal-Wallis test followed by Tukey post hoc correction was used for multiple comparison. For distribution comparisons, a two-sample Kolmogorov-Smirnov test was used. Results were considered significant if p < 0.05.

### Data and Code accessibility

All data generated in this study are included in this article and available upon request. All Matlab scripts used in this study have been deposited at https://github.com/weiwen107/EPhys_analysis_MATLAB.

## Results

### Chronic chemogenetic activity suppression induces synaptic scaling in L2/3 pyramidal neurons during the CP

In order to study the developmental regulation of homeostatic plasticity mechanisms, we needed a paradigm that would allow us to induce comparable activity-deprivation at different developmental stages. DREADD-mediated inhibition *in vivo* has been shown to induce synaptic scaling up in insular cortex (Wu et al., 2021), so we adapted this approach for use in L2/3 pyramidal neurons in V1, where we know that homeostatic and intrinsic plasticity are co-induced by visual deprivation during the classic visual system CP (Lambo and Turrigiano, 2013).

We began by validating the effectiveness of the inhibitory DREADD, hM4Di, under our experimental conditions. An AAV vector carrying hM4Di was stereotaxically delivered into one hemisphere of mouse V1 between P14 and 16 (Fig. 1A). After 7-10 days to allow the virus to express, we observed intense expression of hM4Di in L2/3 pyramidal neurons (Fig. 1B). To confirm that these exogenously expressed hM4Di receptors can be activated by CNO and lead to reduced neuronal firing, we used an active slice preparation to record spontaneous firing in L2/3 pyramidal neurons before and after perfusion with active ACSF containing 500 nM CNO. As expected, CNO wash-in triggered a dramatic reduction in firing rate and a mild hyperpolarization in hM4Di-positive neurons from the infected hemisphere, whereas neurons from the uninfected control hemisphere were not affected by CNO wash-in (Fig.1C).

**Figure 1.**
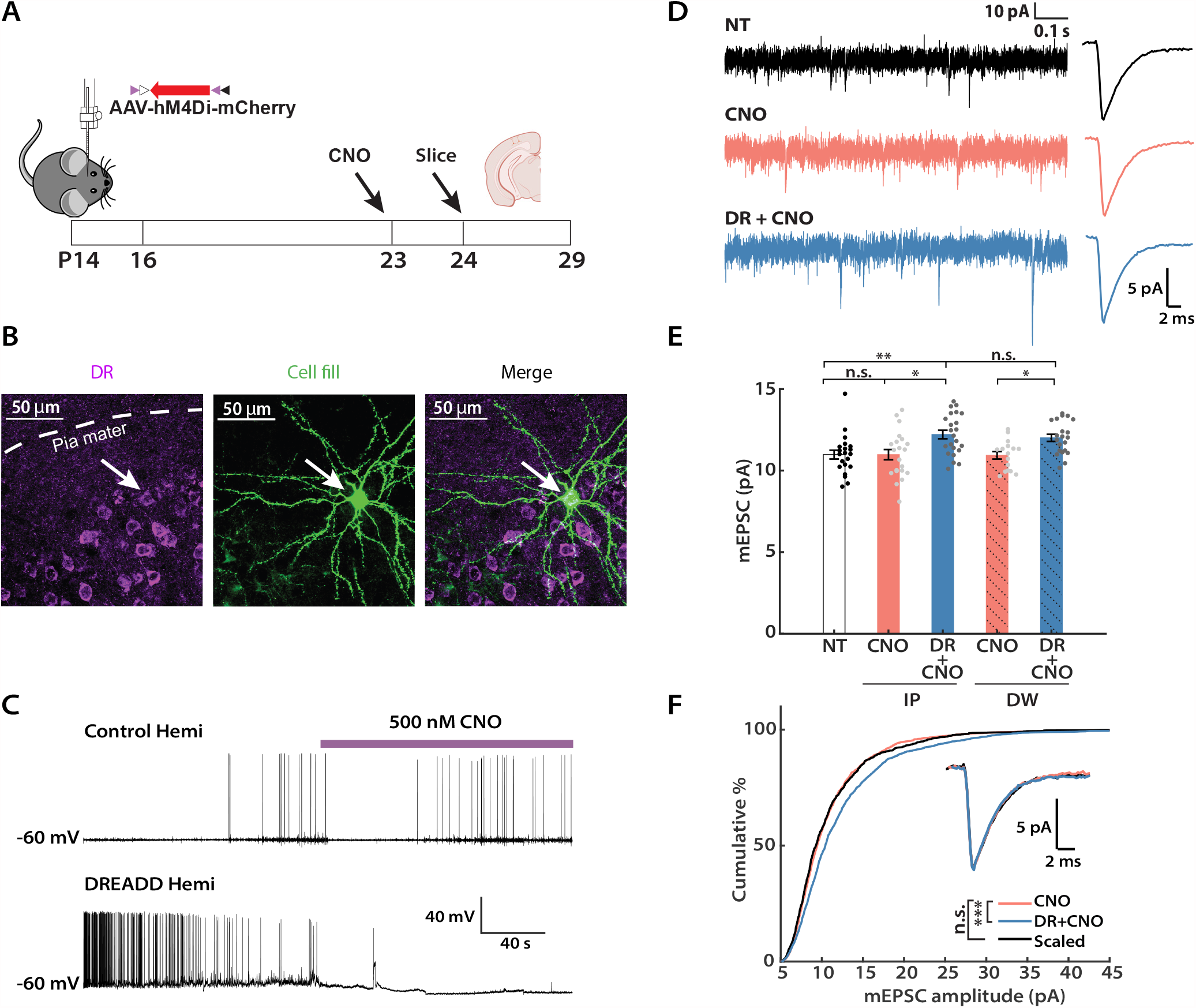
Chronic activity suppression by the inhibitory DREADD induces synaptic scaling up during the classical visual system critical period (CP). **A**, Experimental timeline. Viruses carrying hM4Di were stereotaxically delivered into V1 between P14 and P16. CNO was administered by intraperitoneal injections (IP) or orally via drinking water (DW) for 24 prior to sacrifice. Slice electrophysiology was performed between P24 and P29. Mouse and brain slice icons were adapted from BioRender.com. **B**, Representative images of DREADD expression and biocytin cell fill in L2/3 of mouse V1. Left, DREADD-positive neurons (magenta); middle, a cell filled with biocytin during whole-cell recording (green); right, colocalization of DREADD and biocytin signals. **C**, Validation of hM4Di function via acute CNO perfusion. Representative current clamp recordings of spontaneous firing from the uninfected hemisphere (control hemi) and the infected hemisphere (DREADD hemi), respectively, before and after CNO wash-in. Note the decrease in firing rate and hyperpolarization in the DREADD-positive cell after CNO onset. **D**, Representative raw mEPSC traces and corresponding waveform averages under different conditions. NT: no treatment; CNO: uninfected hemisphere from animals that received CNO; DR + CNO: the DREADD-expressing hemisphere from the same animal. **E**, Comparison of the mean mEPSC amplitude following two different methods of CNO delivery: intraperitoneal injection (IP, solid color) or drinking water (DW, solid color + shadow). NT and two conditions from each delivery method were considered as a group, and Kruskal-Wallis test was performed separately on each group, followed by Tukey post hoc correction (IP, NT vs. CNO: p = 0.9653, NT vs. DR + CNO: p = 0.0069, CNO vs. DR + CNO: p = 0.0154; DW, NT vs. CNO: p = 0.9942, NT vs. DR + CNO: p = 0.0084, CNO vs. DR + CNO: p = 0.0214). An unpaired t test was performed on the two DR + CNO groups (p = 0.1441). **F**, Cumulative distributions of mEPSC amplitudes from the CNO and DR + CNO condition, respectively. Events from the DR + CNO condition were scaled according to the linear function y = 1.26x - 1.30 (black trace). Note that events from these two conditions belong to distinct distributions before scaling but are not significantly different after scaling (Kolmogorov-Smirnov test: CNO vs. DR + CNO: p = 2.50E-12; CNO vs. Scaled: p = 0.1550). Inset: Overlay of peak-scaled waveform average traces from all three conditions, to illustrate mEPSC kinetics. **Sample sizes**: NT, n = 21, 5 animals; IP-CNO, n = 21, 7 animals; IP-DR + CNO, n = 23, 7 animals; DW-CNO, n = 16, 5 animals; DW-DR + CNO, n = 21, 5 animals.

We next designed an experimental paradigm to perturb neuronal activity *in vivo* (Fig. 1A). Inhibitory DREADDs were expressed in V1 as above, and mice received CNO for 24 hours prior to slice electrophysiology. Slice recordings were obtained between P24-P29, well within the classical visual system CP (Espinosa and Stryker, 2012). We compared two methods of CNO delivery: intraperitoneal injections (IP, 5 mg/kg, every 12 hours) as described previously (Wu et al., 2021), and by inclusion of CNO in the drinking water (DW, 0.05 mg/mL, *ad libitum*). Under both delivery methods, the average mEPSC amplitude of hM4Di-positive L2/3 pyramidal neurons (DR + CNO) increased significantly relative to hM4Di-negative neurons from the uninfected hemisphere (Fig. 1D,E. IP: 111.3 ± 3.7%; DW: 109.8 ± 2.9% of control). Importantly, there was no difference between the no-treatment group (NT, representing neurons from mice that experienced neither virus injection nor CNO administration) and the CNO group (neurons from the uninfected hemisphere of mice that received CNO), indicating that CNO administration itself did not affect mEPSC amplitude. Furthermore, when we examined the cumulative distribution function (CDF) of mEPSC amplitudes for the CNO and DR + CNO group, respectively, the latter showed a rightward shift toward larger amplitudes (Fig. 1F, blue vs. orange). When scaled down by a multiplicative factor (Turrigiano et al., 1998; Torrado Pacheco et al., 2021), this distribution was indistinguishable from the CNO alone distribution (Fig. 1F, black vs. orange). No significant changes were observed in mEPSC frequency, kinetics, or passive properties of the recorded neurons across any conditions (Table 1). Because administration of CNO in drinking water is less invasive, we used this delivery method for all subsequent experiments unless otherwise indicated. In summary, these data demonstrate that 24 hours of activity suppression by *in vivo* hM4Di activation can robustly induce synaptic scaling up in L2/3 pyramidal neurons during the CP.

**Table 1.** Passive neuronal properties and kinetics for mEPSC experiments. Fig. 1: Multigroup comparison (NT, IP: CNO, IP: DR + CNO), frequency, p = 0.054; input resistance (R_in_), p = 0.2016; resting membrane potential (V_m_), p = 0.4854; rise time, p = 0.8648; decay τ, p = 0.1773. Two-group comparison (DW: CNO, DW: DR + CNO), frequency, p = 0.2366; R_in_, p = 0.9982; V_m_, p = 0.5476; rise time, p = 0.4422; decay τ, p = 0.7535. Fig. 2: Multigroup comparison (CT+, CT-, GFP), frequency, p = 0.0510; R_in_, p = 0.3211; V_m_, p = 0.1993; rise time, p = 0.3189; decay τ, p = 0.2015. Fig. 4: Multigroup comparison (Shk3 KO: NT, Shk3 KO: CNO, Shk3 KO: DR + CNO), frequency, p = 0.4776; R_in_, p = 0.345; V_m_, p = 0.1637; rise time, p = 0.1724; decay τ, p = 0.8728. Fig. 6: Multigroup comparison (NT, 24h, 48h), frequency, p = 0.053; R_in_, p = 0.1849; V_m_, p = 0.1780; rise time, p = 0.0615; decay τ, p = 0.057.

| Experimental Condition | Frequency (Hz) | $R_{in}$ (MΩ) | $V_m$ (mV) | Rise Time (ms) | Decay $\tau$ (ms) |
| --- | --- | --- | --- | --- | --- |
| Fig. 1 |  |  |  |  |  |
| NT | $4.13 \pm 0.45$ | $151 \pm 7$ | $-74 \pm 2$ | $0.56 \pm 0.02$ | $2.97 \pm 0.12$ |
| IP: CNO | $3.32 \pm 0.47$ | $172 \pm 12$ | $-77 \pm 2$ | $0.56 \pm 0.01$ | $2.72 \pm 0.11$ |
| IP: DR + CNO | $4.59 \pm 0.41$ | $176 \pm 10$ | $-73 \pm 2$ | $0.56 \pm 0.02$ | $2.72 \pm 0.09$ |
| DW: CNO | $4.13 \pm 0.45$ | $189 \pm 7$ | $-81 \pm 1$ | $0.53 \pm 0.01$ | $2.51 \pm 0.07$ |
| DW: DR + CNO | $4.19 \pm 0.46$ | $188 \pm 7$ | $-79 \pm 2$ | $0.54 \pm 0.01$ | $2.42 \pm 0.06$ |
| Fig. 2 |  |  |  |  |  |
| CT+ | $1.54 \pm 0.18$ | $184 \pm 10$ | $-79 \pm 1$ | $0.57 \pm 0.02$ | $3.26 \pm 0.16$ |
| CT- | $2.33 \pm 0.24$ | $202 \pm 7$ | $-82 \pm 1$ | $0.56 \pm 0.01$ | $2.79 \pm 0.10$ |
| GFP | $1.99 \pm 0.28$ | $171 \pm 8$ | $-81 \pm 1$ | $0.59 \pm 0.02$ | $3.00 \pm 0.11$ |
| Fig. 4 |  |  |  |  |  |
| Shk3 KO: NT | $4.09 \pm 0.41$ | $192 \pm 9$ | $-79 \pm 2$ | $0.53 \pm 0.01$ | $2.59 \pm 0.08$ |
| Shk3 KO: CNO | $4.30 \pm 0.37$ | $206 \pm 6$ | $-83 \pm 1$ | $0.51 \pm 0.01$ | $2.51 \pm 0.06$ |
| Shk3 KO: DR + CNO | $4.95 \pm 0.47$ | $209 \pm 10$ | $-82 \pm 2$ | $0.56 \pm 0.01$ | $2.60 \pm 0.08$ |
| Fig. 6 |  |  |  |  |  |
| NT | $2.98 \pm 0.25$ | $146 \pm 7$ | $-80 \pm 2$ | $0.57 \pm 0.01$ | $3.09 \pm 0.08$ |
| 24h | $3.37 \pm 0.27$ | $152 \pm 8$ | $-78 \pm 2$ | $0.59 \pm 0.01$ | $3.04 \pm 0.07$ |
| 48h | $1.90 \pm 0.10$ | $157 \pm 8$ | $-83 \pm 1$ | $0.56 \pm 0.02$ | $2.80 \pm 0.10$ |

### The GluA2 C-tail is required for DREADD-induced synaptic scaling

One of the molecular signatures of deprivation-induced synaptic scaling in neocortical neurons is a reliance on protein interactions involving the GluA2 subunit of the AMPA receptors (Wierenga et al., 2005; Gainey et al., 2009; Goold and Nicoll, 2010). Synaptic scaling induced by TTX in culture and visual deprivation in V1 can be blocked by the expression of GluA2 C-tail fragments (Gainey et al., 2009; Lambo and Turrigiano, 2013). To verify that DREADD-induced scaling in L2/3 neurons operates through the same molecular pathways as classical synaptic scaling, we expressed hM4Di and GluA2 C-tail together (Fig. 2C, CT+) in the left hemisphere, and hM4Di alone (Fig. 2C, CT-) in the right hemisphere to serve as a within-animal control (Fig. 2A), and recoded mEPSCs after 24 hours of CNO delivery. As an additional control, we co-expressed hM4Di and GFP in a vector that shares the same backbone as the C-tail-carrying AAV construct, in a different group of mice (Fig. 2C, GFP). In line with our expectation, neurons that expressed both hM4Di and GluA2 C-tail showed similar mean mEPSC amplitudes (Fig. 2D) to the NT and CNO group (Fig. 1E) following 24 hours of CNO treatment, indicating that these neurons failed to engage synaptic scaling to combat the DREADD-induced chronic silencing. Concomitantly, we observed normal scaling up of mEPSC amplitudes in both the CT- and GFP groups (Fig. 2D. CT-: 113.0 ± 3.2 %; GFP: 110.0 ± 3.6 % of control). We further compared the CDFs of mEPSC amplitude for the CT+ and CT-group. The distribution of the CT-group was significantly shifted to the right compared to the CT+ group (Fig. 2E, pink vs. blue), but once scaled down, the distributions were indistinguishable (Fig. 2E, pink vs. black). There were no significant differences in mEPSC frequency across these three groups (Table 1), and no substantial differences in input resistance, resting membrane potential, or mEPSC kinetics (Table 1). In conclusion, the DREADD-induced global increase in synaptic strength requires GluA2 C-tail interactions, clearly exhibiting a hallmark characteristic of synaptic scaling.

**Figure 2.**
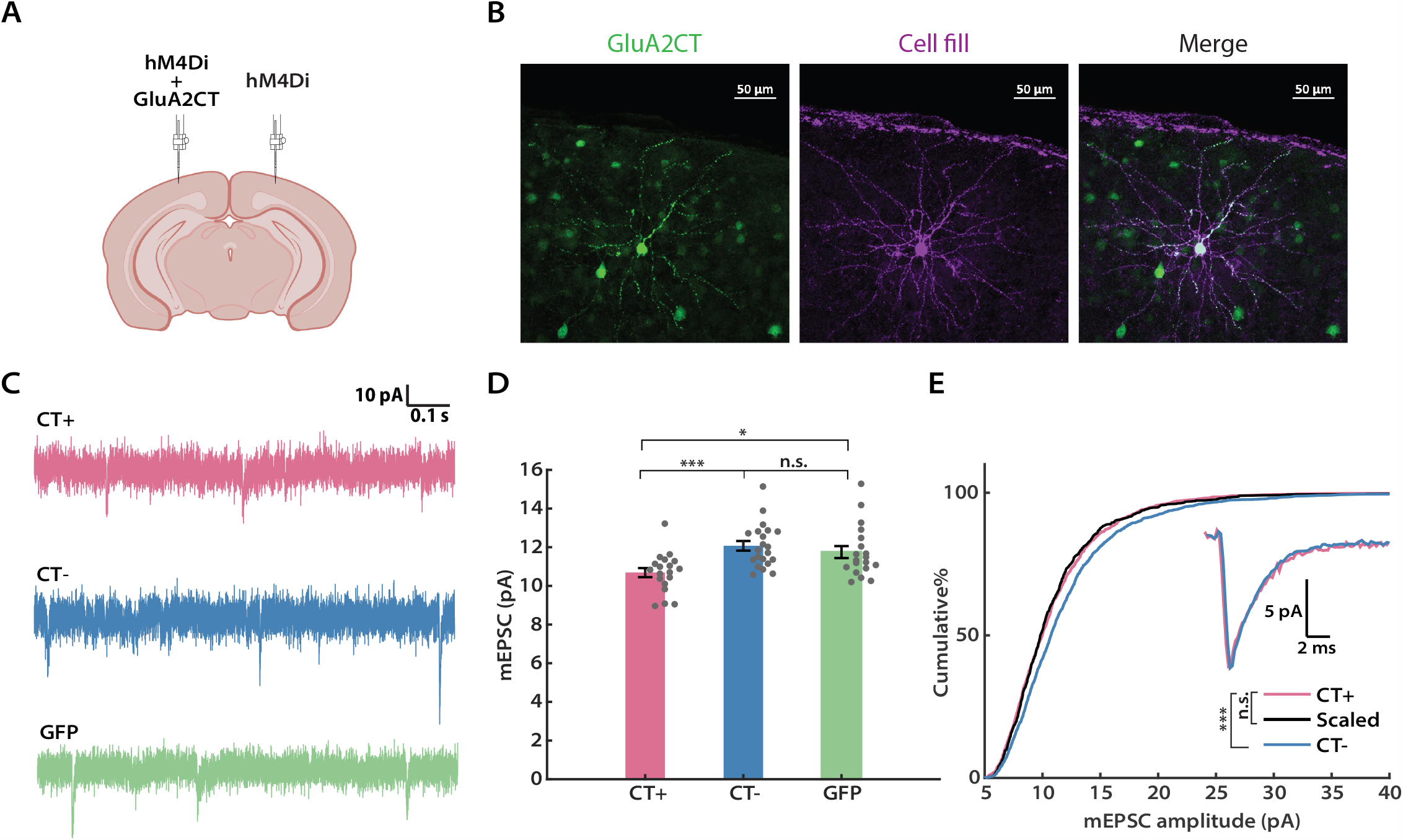
DREADD-induced synaptic scaling is blocked by the GluA2 C-tail. **A**, A schematic of the experimental design (Adapted from icons on BioRender.com). Viruses expressing hM4Di and the GluA2 C-tail, or hM4Di alone, were delivered to the left and right hemispheres of V1, respectively. **B**, Representative images of GluA2 C-tail expression and biocytin cell fill from a L2/3 pyramidal neuron. **C**, Representation mEPSC traces from hM4Di-positive neurons after CNO administration, ± the GluA2 CT (CT+ and CT-), or GFP. **D**, Comparison of the mean mEPSC amplitude from each condition (Kruskal-Wallis test followed by Tukey post hoc correction, CT+ vs. CT-: p = 0.0008; CT+ vs. GFP: p = 0.0388; CT-vs. GFP: p = 0.4924). **E**, Cumulative distributions of mEPSC amplitudes for the indicated conditions. Events from the CT-condition were scaled according to the linear function y = 1.21x -1.10 (black trace). Similarly, two distributions are not significantly different after scaling (Kolmogorov-Smirnov test: CT+ vs. CT-: p = 5.28E-9; CT+ vs. Scaled: p = 0.4355). Inset: Overlay of peak-scaled waveform average traces from all three conditions, to illustrate mEPSC kinetics. **Sample sizes**: CT+, n = 19, 7 animals; CT-, n = 21, 7 animals; GFP, n = 19, 6 animals.

### DREADD-induced activity suppression increases intrinsic excitability of L2/3 pyramidal neurons during the CP

Following prolonged visual deprivation during the CP, L2/3 pyramidal neurons simultaneously engage synaptic scaling and intrinsic homeostatic plasticity to restore overall activity (Hengen et al., 2013; Lambo and Turrigiano, 2013). We therefore wished to know whether hM4Di silencing *in vivo* during the CP was also able to induce homeostatic changes in intrinsic excitability. To investigate this, we expressed hM4Di as described above, administered CNO for 24 hours (via IP injection), then cut acute slices and obtained whole-cell current clamp recordings from L2/3 pyramidal neurons, and generated f-I curves in the presence of synaptic blockers (Fig. 3A). Indeed, we found that the hM4Di-positive neurons exhibited an upward- and leftward-shifted f-I curve, characterized by a higher mean instantaneous firing rate (mean IFR) at all current steps (Fig. 3B, left panel). Quantification of the area under the f-I curve (Trojanowski et al., 2021) for each neuron demonstrated that this measure increased significantly following hM4Di-mediated inhibition (158.1 ± 14.4 % of control), while CNO alone had no impact (Fig. 3B, right panel). To dissect the underlying cellular properties that could lead to such an increase, we further analyzed the firing pattern and the action potential shape of neurons from each group, respectively. In line with a higher intrinsic excitability, when compared with neurons from the other two groups, hM4Di-positive neurons exhibited a lower rheobase (Fig. 3C. 81.1 ± 7.8 % of control), less spike frequency adaptation (Fig. 3D. 56.5 ± 16.4 % of control), and shorter latency to the first spike (Fig. 3E. 75.0 ± 6.3 % of control). In contrast, we did not see any change in the threshold voltage, the amplitude of the afterhyperpolarization, or the action potential width at half-maximum (Fig. 3G,H,I). We further generated the waveform average trace of the first action potentials evoked at the rheobase for each group. Consistent with above findings, action potential waveforms from CNO and DR + CNO groups were indistinguishable (Fig. 3F). Under our mEPSC recording conditions, input resistance was slightly but not significantly increased by activity-suppression (Table 1); however, under these f-I recording conditions input resistance did increase significantly (Table 2), similar to our previous observations following monocular deprivation (Lambo and Turrigiano, 2013). Together, these results demonstrate that the two major forms of homeostatic plasticity, synaptic scaling and intrinsic homeostatic plasticity, work in parallel to restore neuronal excitability of L2/3 pyramidal neurons following hM4Di-induced activity suppression during the CP.

**Table 2.** Passive neuronal properties for intrinsic excitability experiments Fig. 3: multigroup comparison (NT, CNO, DR + CNO), R_in_, NT vs. CNO, p = 0.7491, NT vs. DR + CNO, p = 0.0032, CNO vs. DR + CNO, p = 0.0234; V_m_, p = 0.7837. Fig. 5: two-group comparison, R_in_, p = 0.4647; V_m_, p = 0.3315. Fig. 6: multigroup comparison (CNO, DR + 24h, DR + 48h), R_in_, CNO vs. DR + 24h, p = 0.0372, CNO vs. DR + 48h, p=0.0290, DR + 24h vs. DR + 48h, p = 0.9966; V_m_, p = 0.3465.

| Experimental | $R_{in}$ | $V_m$ |
| --- | --- | --- |
| Condition | (MΩ) | (mV) |
| Fig. 3 |  |  |
| NT | $141 \pm 6$ | $-76 \pm 1$ |
| CNO | $148 \pm 7$ | $-75 \pm 1$ |
| DR + CNO | $181 \pm 10$ | $-74 \pm 1$ |
| Fig. 5 |  |  |
| Shk3 KO: CNO | $193 \pm 12$ | $-80 \pm 1$ |
| Shk3 KO: DR + CNO | $203 \pm 10$ | $-82 \pm 1$ |
| Fig. 7 |  |  |
| CNO (pooled control) | $143 \pm 6$ | $-82 \pm 1$ |
| DR + 24h | $165 \pm 10$ | $-80 \pm 1$ |
| DR + 48h | $162 \pm 10$ | $-82 \pm 1$ |

**Figure 3.**
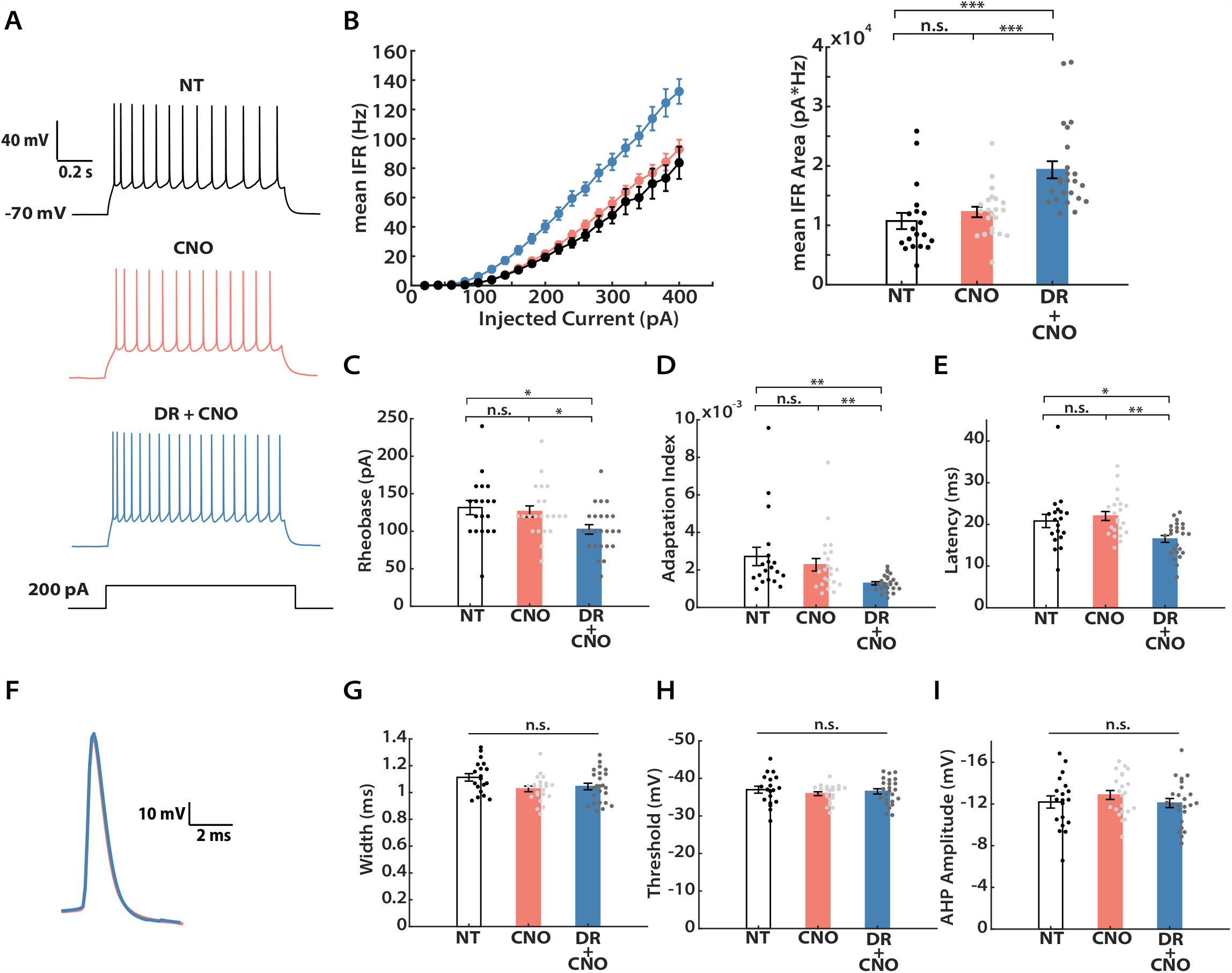
DREADD-induced chronic activity suppression increases intrinsic excitability. **A**, Representative recordings from L2/3 pyramidal neurons under three conditions (NT, CNO, DR + CNO) evoked by a 200 pA current injection. **B**, Left: frequency vs. current (f-I) curves for the three conditions. Y axis indicates mean instantaneous firing rate (IFR). Right: quantification of the area under each f-I curve, calculated individually for each neuron and then averaged for each condition. A Kruskal-Wallis test followed by Tukey post hoc correction was performed (NT vs. CNO: p = 0.2063; NT vs. DR + CNO: p = 1E-6; CNO vs. DR + CNO: p = 1E-4). **C**, Mean rheobase for each condition (Kruskal-Wallis test followed by Tukey post hoc correction: NT vs. CNO: p = 0.8902; NT vs. DR + CNO: p = 0.0103; CNO vs. DR + CNO: p = 0.0306). **D**, Mean adaptation index (Kruskal-Wallis test followed by Tukey post hoc correction: NT vs. CNO: p = 0.7401; NT vs. DR + CNO: p = 0.0011; CNO vs. DR + CNO: p = 0.0099). **E**, Mean latency to the first spike for 300 pA current injection (Kruskal-Wallis test followed by Tukey post hoc correction: NT vs. CNO: p = 0.6692; NT vs. DR + CNO: p = 0.0362; CNO vs. DR + CNO: p = 0.0015). **F**, Overlay of waveform average trace for the first evoked action potentials at rheobase, from CNO and DR + CNO conditions, indicating no change in spike waveform. **G**, Spike widths at half maximum height (Kruskal-Wallis test followed by Tukey post hoc correction, p = 0.0651). **H**, Action potential voltage thresholds (Kruskal-Wallis test followed by Tukey post hoc correction, p = 0.6681). **I**, Afterhyperpolarization (AHP) amplitudes (Kruskal-Wallis test followed by Tukey post hoc correction, p = 0.5062). **Sample sizes**: NT, n = 19, 3 animals; CNO, n = 22, 7 animals, DR + CNO, n = 24, 7 animals.

### Shank3 Knockout mice show deficits in DREADD-induced homeostatic plasticity

We recently reported that acute knockdown of Shank3 in young cultured neocortical neurons results in simultaneous loss of synaptic scaling and intrinsic homeostatic plasticity, as well as loss of firing rate homeostasis in Shank3 KO mice (Tatavarty et al., 2020). We therefore wished to assess whether the hM4Di-mediated induction of synaptic scaling and intrinsic homeostatic plasticity are absent in these mice. To examine this possibility, we repeated the *in vivo* hM4Di silencing paradigm in Shank3B^-/-^ KO animals (Peça et al., 2011; Tatavarty et al., 2020) and wild-type littermates (WT). We firstly compared the ability of the two groups to express synaptic scaling during the CP (Fig. 4A). CNO was administered via drinking water, and the average CNO consumption did not differ between the two groups (WT, 0.28 ± 0.01 mg; Shank3 KO, 0.29 ± 0.03 mg; unpaired t test: p = 0.8067). Analyses of the mEPSC recordings revealed that while mEPSC amplitudes of L2/3 pyramidal neurons were scaled up as expected in WT littermates (Fig. 4B, left three columns; 4D), there was no significant increase in Shank3 KO animals (Fig. 4B, right three columns, 4E). There were no significant differences in mEPSC frequency, kinetics, or passive neuronal properties in Shank3 KO animals (Fig. 4C, Table 1). Therefore, DREADD-induced synaptic scaling is absent in these animals.

**Figure 4.**
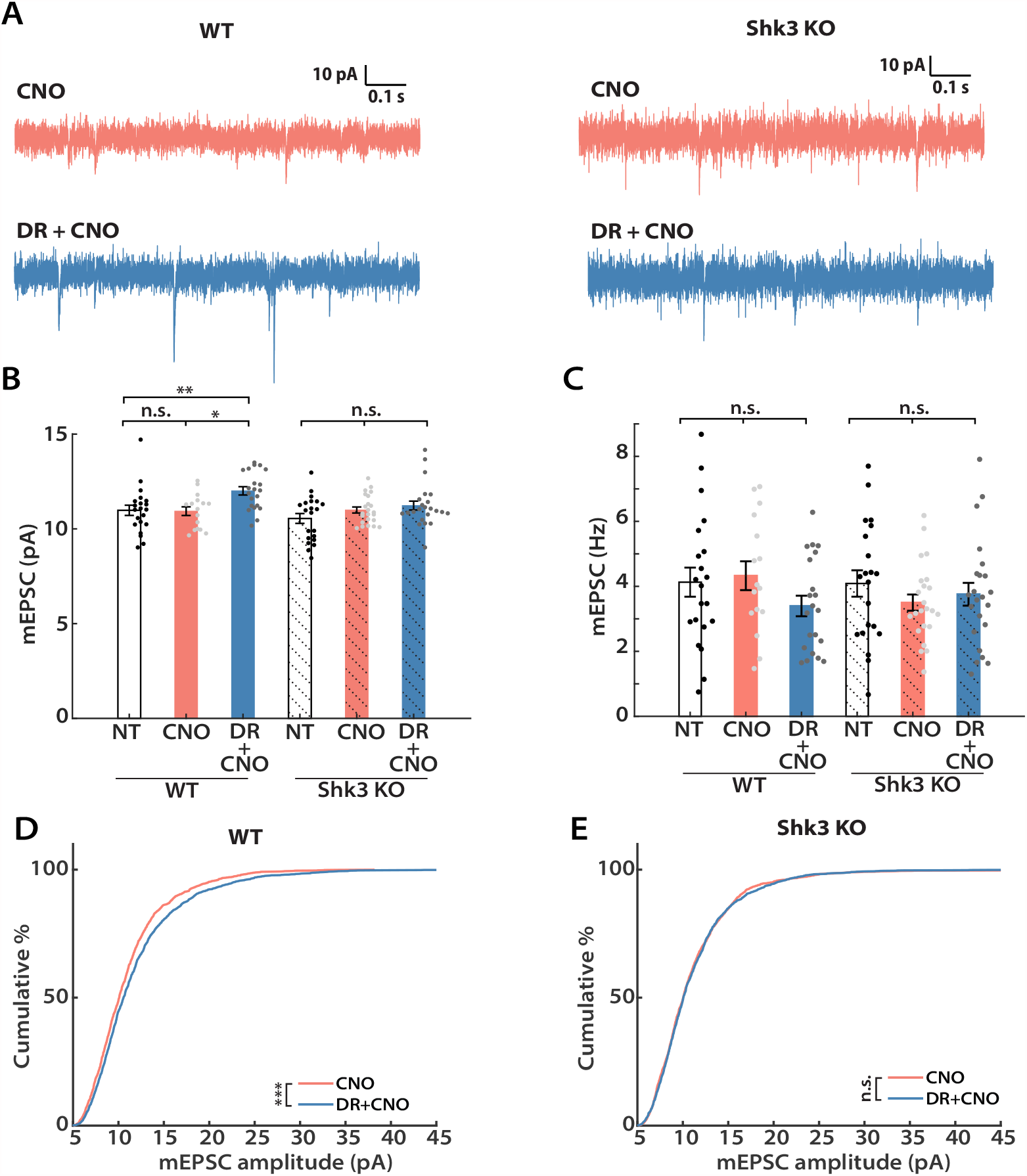
DREADD-induced synaptic scaling is impaired in Shank3 knockout mice. **A**, Representative mEPSC traces and their waveform averages from Shank3 knockouts (Shk3 KO, right) and wild-type littermates (WT, left). **B**, Mean mEPSC amplitude following CNO administration from WT (solid color) and Shk3 KO (solid color + shadow) littermates. Both groups underwent a Kruskal-Wallis test followed by Tukey correction (For WT, NT vs. CNO: p = 0.9942, NT vs. DR + CNO: p = 0.0084, CNO vs. DR + CNO: p = 0.0214; for Shank3 KO, p = 0.3720). **C**, Comparisons of the mean mEPSC frequency from each group (one-way ANOVA followed by Tukey correction: WT, p = 0.2366; Shk3 KO, p = 0.4776). **D**, Cumulative distributions of mEPSC amplitudes from WT littermates (Kolmogorov-Smirnov test: CNO vs. DR + CNO: p = 2.28E-4). **E**, Same as D, but for Shk3 KO animals (Kolmogorov-Smirnov test: CNO vs. DR + CNO: p = 0.5456). **Sample sizes**: WT-NT, n = 21, 5 animals; WT-CNO, n = 16, 5 animals; WT-DR + CNO, n = 21, 5 animals; Shk3 KO-NT, n = 21, 3 animals; Shk3 KO-CNO, n = 23, 6 animals; Shk3 KO-DR + CNO, n = 23, 6 animals.

Next, we assessed intrinsic excitability (Fig. 5). Examination of the f-I curves from WT and KO littermates revealed two clear effects (Fig. 5A, Fig. 5B, left panel). First, KO neurons showed greater intrinsic excitability at baseline than WT neurons. Second, while DREADD-mediated inhibition increased the intrinsic excitability of WT neurons as expected, neurons from the KO group showed no change in intrinsic excitability. A similar pattern was observed for peak instantaneous firing rate (Fig. 5C). Furthermore, KO neurons were similar to the WT DREADD group in possessing lower rheobase (Fig. 5D. 68.6±8.0 % of control), shorter latency to the first spike (Fig. 5E. 67.2 ± 7.1 % of control), and less spike adaptation (Fig. 5F. 63.4 ± 16.6 % of control). Thus, Shank3 loss has rendered these L2/3 pyramidal neurons more excitable, and has possibly occluded the normal activity-dependent change in intrinsic excitability observed in WT animals. In summary, both the global increase in mEPSC amplitudes and the shift in f-I curves that are normally induced by DREADD-mediated inhibition are absent in Shank3 KO animals.

**Figure 5.**
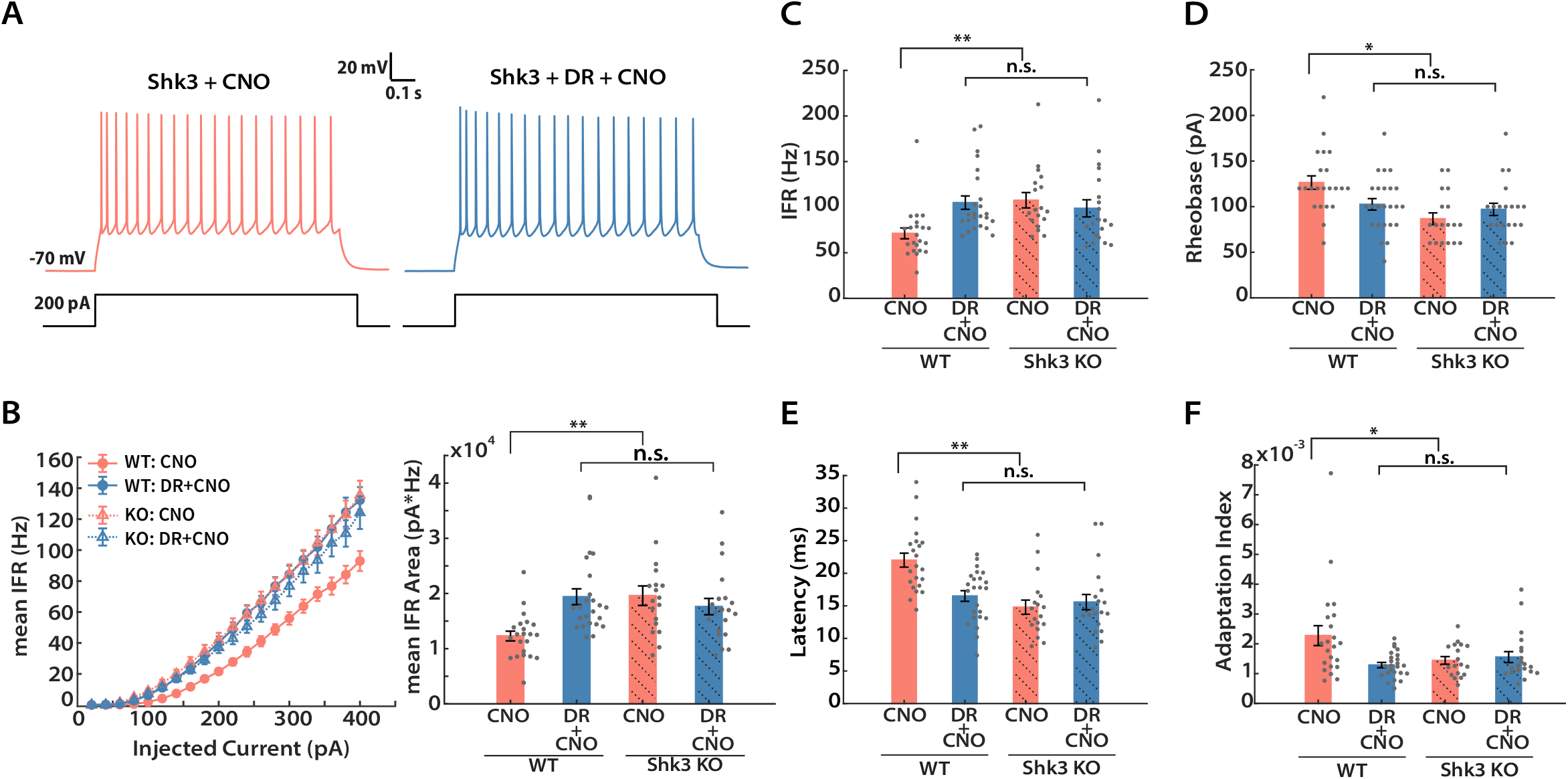
DREADD-induced intrinsic homeostatic plasticity is impaired in Shank3 knockout mice. **A**, Representative recordings from Shk3 KO neurons under the indicated conditions. **B**, Mean IFR from WT (solid circle, solid line) and Shk3 KO (hollow triangle, dashed line) animals under DREADD manipulation (left) and quantification of areas under the curve (right). Kruskal-Wallis test followed by Tukey correction was performed (WT vs. WT + DR, p = 0.0003; WT vs. Shk3, p = 0.0004; WT vs. Shk3 + DR, p = 0.0113; WT + DR vs. Shk3, p = 0.9934; WT + DR vs. Shk3 + DR, p = 0.8657; Shk3 vs. Shk3 + DR, p = 0.7619). **C**, Fraction of neurons with IFR > 100 Hz (at 300 pA current injection) (Kruskal-Wallis test followed by Tukey post hoc correction: WT vs. WT + DR, p = 0.0008; WT vs. Shk3, p = 0.0006; WT vs. Shk3 + DR, p = 0.0458; WT + DR vs. Shk3, p = 0.9875; WT + DR vs. Shk3 + DR, p = 0.7086; Shk3 vs. Shk3 + DR, p = 0.5544). **D**, Mean rheobase (Kruskal-Wallis test followed by Tukey post hoc correction: WT vs. WT + DR, p = 0.011; WT vs. Shk3, p = 0.0008; WT vs. Shk3 + DR, p = 0.0184; WT + DR vs. Shk3, p = 0.2951; WT + DR vs. Shk3 + DR, p = 0.8609; Shk3 vs. Shk3 + DR, p = 0.7765). **E**, Mean latency to first spike at 300 pA current injection (Kruskal-Wallis test followed by Tukey post hoc correction: WT vs. WT + DR, p = 0.01; WT vs. Shk3, p = 0.0001; WT vs. Shk3 + DR, p = 0.0004; WT + DR vs. Shk3, p = 0.4851; WT + DR vs. Shk3 + DR, p = 0.7495; Shk3 vs. Shk3 + DR, p = 0.9727). **F**, Mean adaptation index (at 300 pA current injection) (Kruskal-Wallis test followed by Tukey post hoc correction: WT vs. WT + DR, p = 0.0244; WT vs. Shk3, p = 0.0318; WT vs. Shk3 + DR, p = 0.0205; WT + DR vs. Shk3, p = 9187; WT + DR vs. Shk3 + DR, p = 0.8727; Shk3 vs. Shk3 + DR, p = 0.9997). **Sample sizes**: WT-CNO and -DR + CNO, same as indicated in Figure 3; Shk3 KO-CNO, n = 20, 4 animals; Shk3 KO-DR + CNO, n = 18, 4 animals.

### Developmental regulation of intrinsic homeostatic plasticity

To determine whether homeostatic plasticity is maintained into adulthood, we used the same DREADD-induced inhibition paradigm in adult animals. We again administered CNO for 24 hours, then performed slice electrophysiology experiments between P45 and P55, well after the end of the classical rodent visual system CP (Espinosa and Stryker, 2012). Interestingly, we still observed a significant increase in the mean mEPSC amplitude after 24 hours of CNO administration (Fig. 6A,B. 24h, 110.1 ± 2.7% of control). We next wondered if longer deprivation would induce stronger synaptic scaling in adult animals as it does during the CP (Lambo and Turrigiano, 2013). To investigate this possibility, we repeated these mEPSC measurements after an additional 24 hours of activity suppression. Longer deprivation indeed induced a further increase in the mean amplitude (Fig. 6A,B. 48h, 117.7 ± 3.0% of control). Similar to their younger counterparts, passive neuronal properties and mEPSC kinetics were not significantly different across conditions (Table 1). Thus, robust synaptic scaling is still present in adult V1.

**Figure 6.**
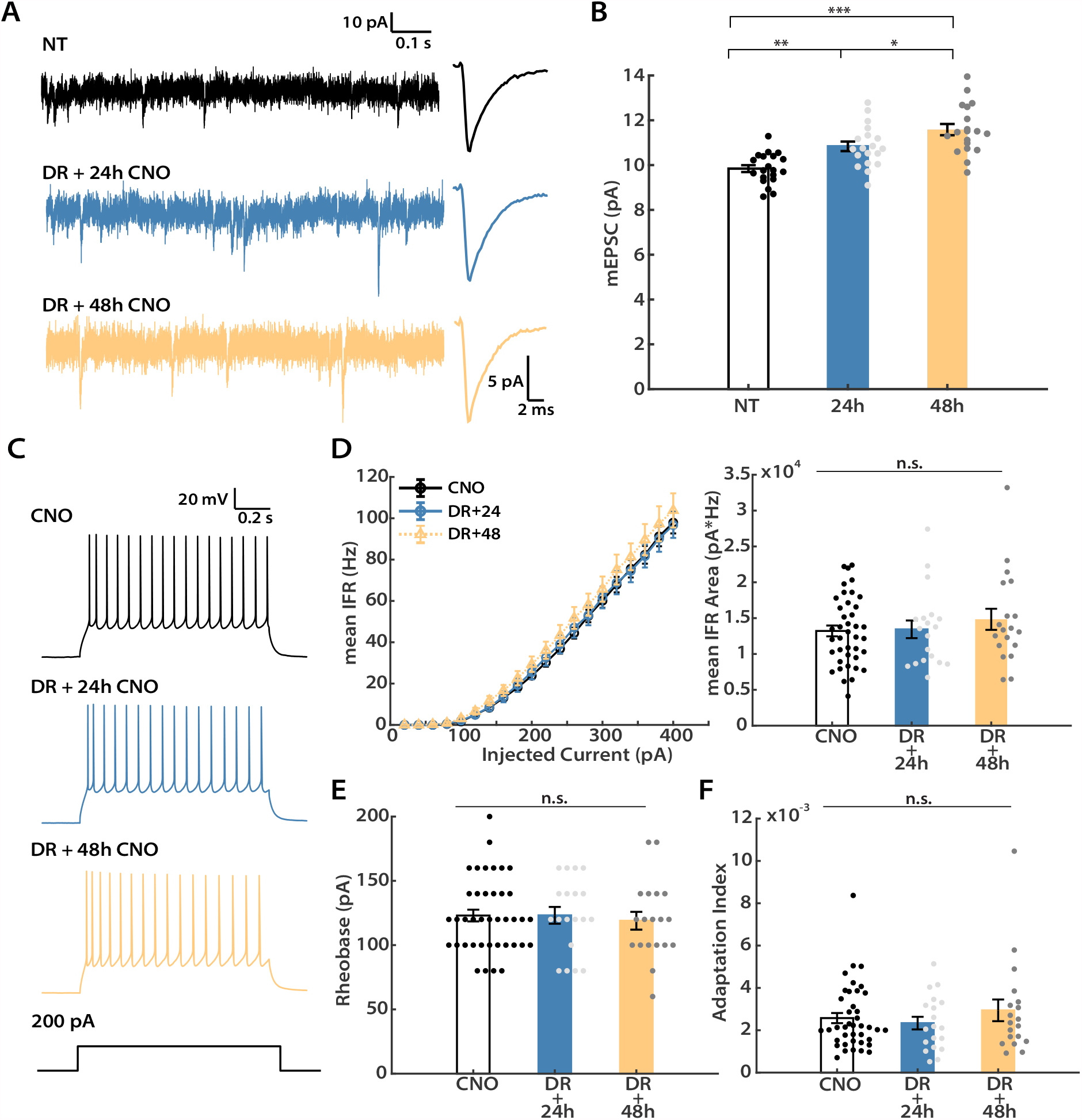
DREADD-induced synaptic scaling persists into adulthood, while intrinsic homeostatic plasticity does not. Recordings were performed on adult mice (P45-55) after hM4Di expression in one hemisphere, and either 24 or 48 hours of CNO administration. **A**, Representative mEPSC traces and corresponding waveform averages from three conditions: NT (no treatment), DR + 24h (24 hours of CNO), DR + 48h (48 hours of CNO). **B**, Mean mEPSC amplitude for the three conditions (one-way ANOVA followed by Tukey post hoc correction: NT vs. 24h, p = 0.0036; NT vs. 48h, p = 6.0E-7; 24h vs. 48h, p = 0.0393). **C**, Representative evoked firings from all three conditions. **D**, Left: F-I curves generated from neurons under each condition. Y-axis indicates the mean IFR. Right: Quantification of the area under each f-I curve (Kruskal-Wallis test followed by Tukey post hoc correction: p = 0.7341). Data from the uninfected hemisphere after 24h and 48h CNO were not significantly different and were pooled for the CNO condition. **E**, Mean rheobase (Kruskal-Wallis test followed by Tukey post hoc correction: p = 0.8205). **F**, mean adaptation index evoked by 300 pA current injection (Kruskal-Wallis test followed by Tukey post hoc correction: p = 0.8148). **Sample sizes for mEPSC experiments**: CNO, n = 39, 10 animals, DR + 24h, n = 19, 6 animals, DR + 48h, n = 19, 4 animals. **Sample sizes for intrinsic excitability experiments**: NT, n = 20, 3 animals; DR + 24h, n = 19, 6 animals, DR + 48h, n = 19, 4 animals.

Next, we probed for intrinsic homeostatic plasticity in adult animals (Fig. 6C). To our great surprise, there was no discernable change in intrinsic excitability of hM4Di-positive neurons following either 24 or 48 hours of activity suppression, as indicated by both the almost superimposable f-I curves (Fig. 6D, left panel), and the lack of change in mean area under the f-I curves (Fig. 6D, right panel). Cell properties that contributed to the elevated intrinsic excitability in juvenile animals, such as rheobase and spike adaptation, were also not different in adult animals at either time point (Fig. 6E,F). Interestingly, consistent with our observation in CP animals, there was also a small (∼13%) increase in input resistance under these recording conditions (Table 2). However, these changes were not sufficient to alter f-I curves or other measures of excitability in adult animals. Taken together, our data show that, while synaptic scaling is intact, intrinsic homeostatic plasticity is absent in L2/3 pyramidal neurons in adult animals.

### Developmental regulation of inhibitory homeostatic plasticity

Plasticity of inhibition can play a prominent role in the adult visual cortex (Kameyama et al., 2010; Ribic, 2020). We therefore wondered if neurons in the adult visual cortex might engage inhibitory homeostatic mechanisms in place of intrinsic homeostatic plasticity. To investigate this possibility, we recorded mIPSCs from both DREADD-positive and control L2/3 pyramidal neurons following 24 hours of DREADD-induced activity-suppression, from both CP and adult animals (Fig. 7A,B). MIPSC were recorded as inward currents from a holding potential of -70 mV in a symmetrical chloride solution. Interestingly, mIPSC amplitude was not affected by activity-suppression at either age (Fig. 7C). In CP animals, mIPSC frequency was also unaffected by activity suppression (Fig. 7D, left two columns). In marked contrast, there was a 52% reduction in mean mIPSC frequency in adult L2/3 pyramidal neurons (Fig. 7C, right two columns), indicating that spontaneous release events are dramatically reduced following chronic activity-suppression. We observed no major changes in passive neuronal properties or mIPSC kinetics for any condition (Table 3). Taken together, the above results show that, following the same activity perturbation, L2/3 pyramidal neurons engage different sets of homeostatic plasticity mechanisms at distinct developmental stages. After CP closure, these neurons no longer homeostatically adjust intrinsic excitability, but instead adjust inhibition.

**Table 3.** Passive neuronal properties and kinetics for mIPSC experiments Two-group comparison (Adult, CP): R_in_, adult, p = 0.054, CP, p = 0.1263; V_m_, adult, p = 0.6647, CP, p = 0.8804; rise time, adult, p = 0.7544, CP, p = 0.4351; decay_fast, adult, p = 0.7637, CP, p = 0.5792; decay_slow, adult, p = 0.0028, CP, p = 0.9031; fast%, adult, p = 0.6554, CP, p = 0.1080.

| Experimental | $R_{in}$ | $V_m$ | Rise Time | Decay_fast | Decay_slow | Fast |
| --- | --- | --- | --- | --- | --- | --- |
| Condition | (MΩ) | (mV) | (ms) | (ms) | (ms) | (%) |
| Fig. 8 |  |  |  |  |  |  |
| Adult: CNO | $171 \pm 8$ | $-81 \pm 1$ | $0.43 \pm 0.01$ | $3.38 \pm 0.18$ | $53.59 \pm 8.09$ | $80.2 \pm 4.3$ |
| Adult: DR + CNO | $152 \pm 8$ | $-81 \pm 1$ | $0.45 \pm 0.02$ | $3.34 \pm 0.16$ | $21.77 \pm 5.30$ | $70.9 \pm 3.7$ |
| CP: CNO | $135 \pm 8$ | $-82 \pm 1$ | $0.44 \pm 0.02$ | $3.61 \pm 0.13$ | $20.80 \pm 2.10$ | $74.5 \pm 3.3$ |
| CP: DR + CNO | $154 \pm 10$ | $-81 \pm 1$ | $0.44 \pm 0.01$ | $3.56 \pm 0.17$ | $27.15 \pm 5.43$ | $76.6 \pm 2.8$ |

**Figure 7.**
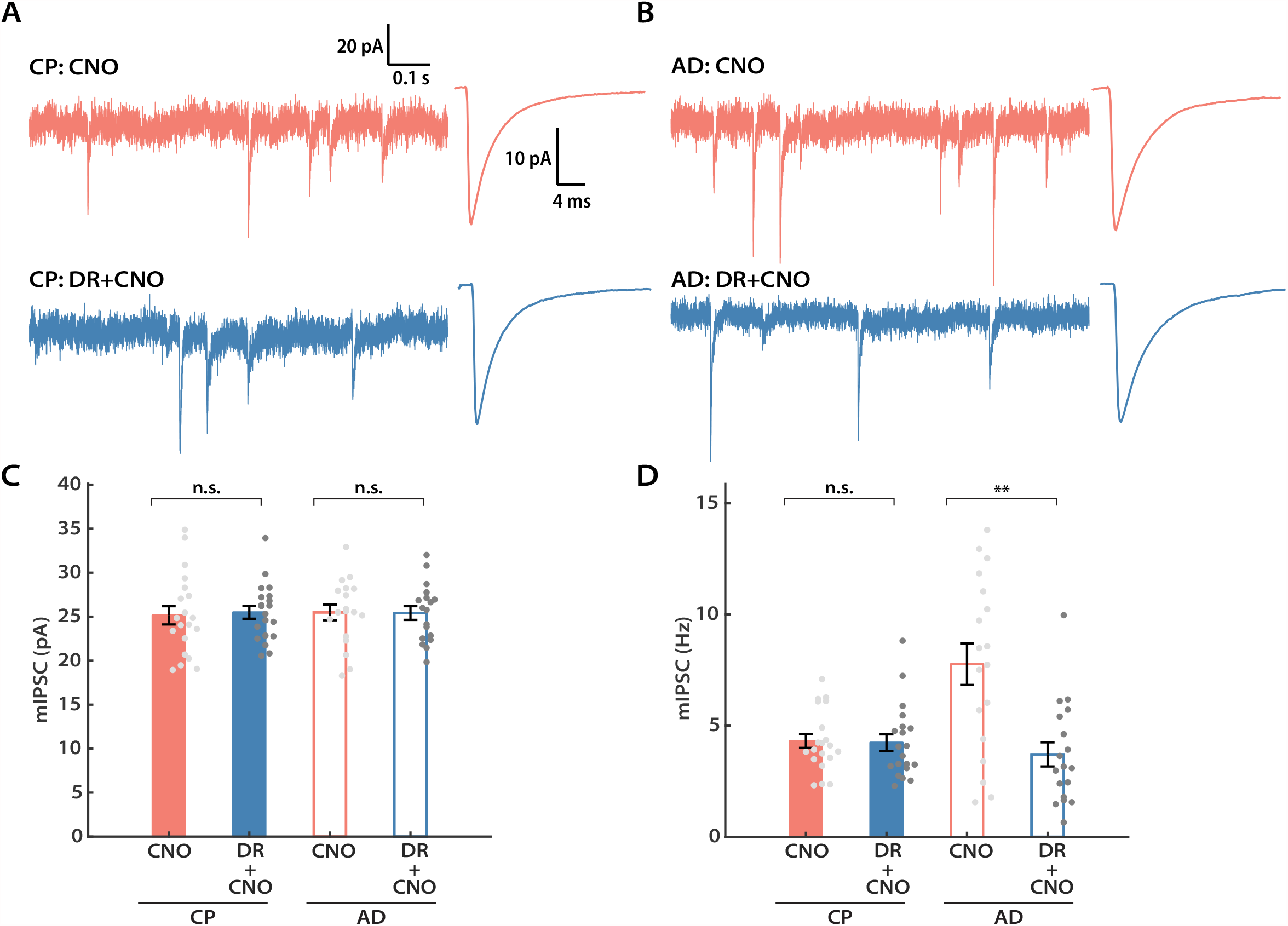
L2/3 pyramidal neurons engage inhibitory plasticity following DREADD-induce chronic silencing in adulthood (AD) but not during the critical period (CP). **A**, Representative mIPSC traces and corresponding waveform averages from the control (CNO) and DREADD (DR + CNO) conditions during CP. **B**, Same as **A**, but for adult animals. **C**, Mean mIPSC amplitude from both conditions for CP and AD, respectively (CP, unpaired t test: p = 0.4351; AD, unpaired t test: p = 0.8868). **D**, Same as **C**, but comparing the mean mIPSC frequency (CP, Mann-Whitney U test: p = 0.5978; AD, Mann-Whitney U test: p = 0.0029). **Sample sizes**: for AD experiments, both conditions, n = 18, 5 animals; for CP experiments, both conditions, n = 20, 5 animals.

## Discussion

While homeostatic compensation has been widely observed within sensory cortex, the developmental regulation of homeostatic plasticity remains poorly understood. Here we developed an *in vivo* approach (DREADD-mediated activity suppression) that allowed us to manipulate activity similarly in young and adult V1. This paradigm induced classical excitatory synaptic scaling up in L2/3 pyramidal neurons, which depended on GluA2-mediated trafficking mechanisms and the synaptic scaffold protein Shank3, as it does *in vitro* (Gainey et al., 2015; Tatavarty et al., 2020). We found that homeostatic compensation within L2/3 pyramidal neurons is dramatically different in CP and adult animals (Fig. 8). While excitatory synaptic scaling and intrinsic homeostatic plasticity cooperate to restore excitability during the CP, intrinsic homeostatic plasticity is absent in adults. Further, plasticity of quantal inhibitory transmission onto L2/3 neurons is absent during the CP, but is robustly recruited in adults. Our data suggest that, rather than being redundant, individual homeostatic mechanisms subserve distinct aspects of excitability maintenance, and can be turned on or off depending on the current needs of neurons and circuits.

**Figure 8.**
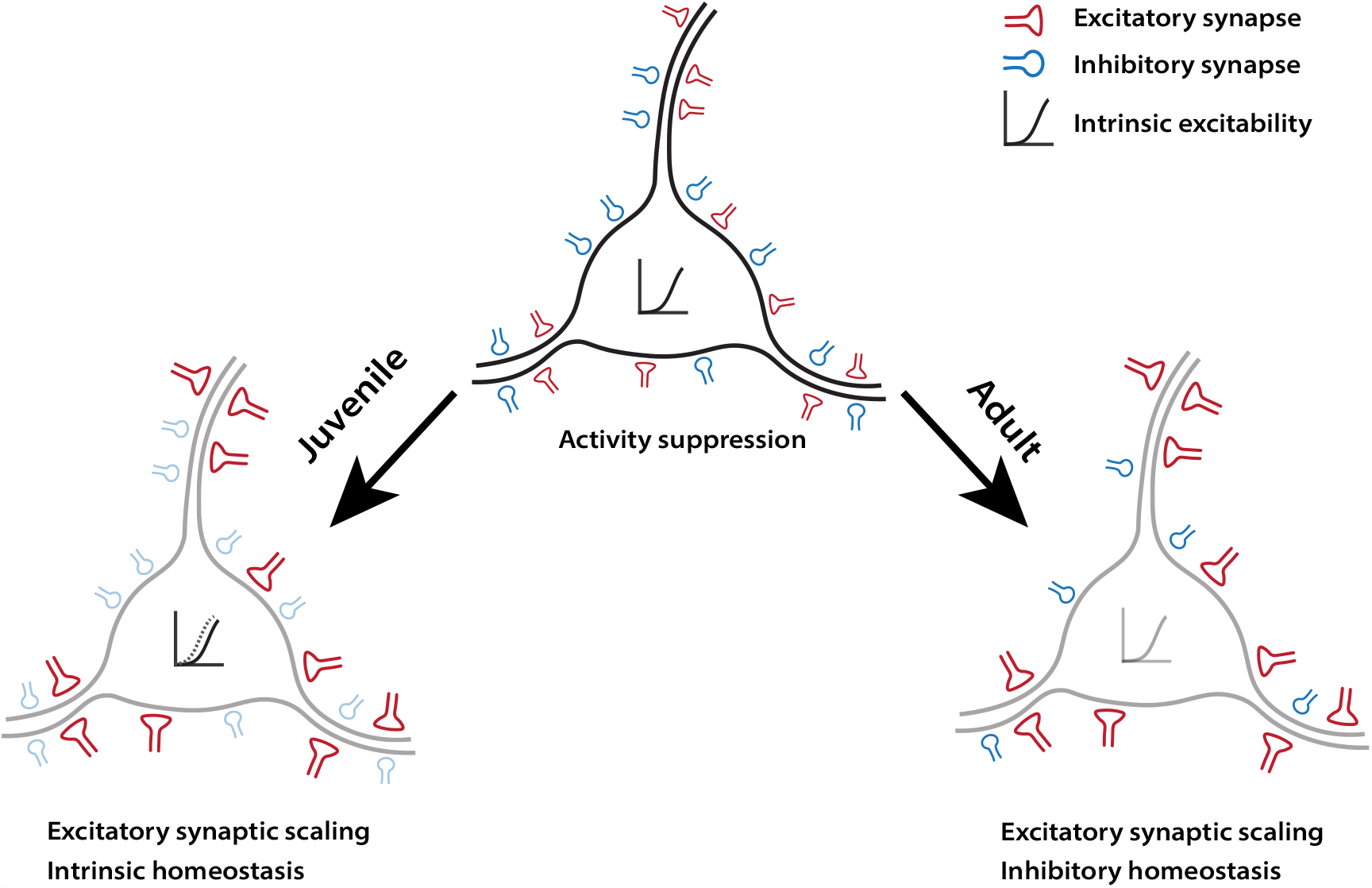
Summary illustration of the distinct homeostatic responses of L2/3 pyramidal neurons during different developmental stages. During the classical visual system CP (Juvenile), neurons engage excitatory synaptic scaling and intrinsic homeostatic plasticity in response to DREADD-induced activity suppression (left). However, while excitatory synaptic scaling persisted into adulthood, intrinsic mechanisms are replaced by inhibitory homeostatic compensations (right).

When compared to other *in vivo* homeostatic plasticity induction methods, DREADDs have several advantages. Most work on neocortical homeostatic plasticity has been limited to sensory cortex following sensory deprivation (Turrigiano, 2011; Gainey and Feldman, 2017). However, the complexity of activity changes during sensory deprivation makes it difficult to disentangle direct from indirect compensations. First, sensory deprivation induces changes along the entire sensory pathway that depend on the deprivation method, and for many paradigms, the induction of homeostatic plasticity requires the prior induction of Hebbian plasticity (Espinosa and Stryker, 2012; Hengen et al., 2016; Torrado Pacheco et al., 2021). Since identical manipulations can produce distinct forms of Hebbian plasticity in juvenile and adult animals (Cooke and Bear, 2014; Hübener and Bonhoeffer, 2014), their impact on neocortical activity at different developmental stages is rarely equivalent, and thus the site(s) and time-course(s) of homeostatic compensation may differ. DREADDs allow us to circumvent both limitations. With the ability to restrict DREADD expression to specific neuronal types in desired regions, and the direct action of DREADDs on these neurons, we can induce comparable activity changes in the same subset of neurons at different developmental timepoints. Second, the timeline of DREADD-induced homeostatic plasticity is significantly shorter. DREADD activation robustly induces homeostatic changes within 24 hours; in contrast, sensory perturbations generally take days to induce commensurate changes (Lambo and Turrigiano, 2013; Greenhill et al., 2015; Teichert et al., 2017), mainly because activity suppression in neocortex itself takes time to develop (Espinosa and Stryker, 2012; Hengen et al., 2013, 2016; Pacheco et al., 2019). Lastly, the flexibility of viral-mediated DREADDs expression allows the direct activity manipulation of targeted cells in non-sensory cortical and subcortical areas (Sternson and Roth, 2014), potentially expanding the scope of homeostatic plasticity research *in vivo*.

There are a number of technical issues to consider when adopting DREADDs for chronic activity manipulations. First, the efficacy of DREADDs depends critically on the local availability of the activating ligand. This is especially a challenge for long-term manipulations in the CNS, where ligand concentration diminishes with time after administration (Guettier et al., 2009); although some behavioral phenotypes can last for as long as 8 hours (Alexander et al., 2009). Drinking water delivery has an advantage over bolus administration for chronic manipulations, as animals have a continuous intake of the ligand, but there may still be variations in ligand concentration depending on when animals drink. Despite these caveats, we found that *in vivo* DREADD activation via both delivery methods induced an increase in mEPSC amplitude that was comparable to that observed following visual deprivation (Desai et al., 2002; Hengen et al., 2013; Lambo and Turrigiano, 2013). Second, off-target effects of CNO or other ligands have been reported after acute CNO application (MacLaren et al., 2016; Jendryka et al., 2019), although CNO concentrations in the cerebrospinal fluid following doses as high as 10 mg/kg are still low enough to keep the off-target effects minimal (Jendryka et al., 2019). Here we found that synaptic and intrinsic properties from our within-animal controls from the uninfected hemisphere (exposed to CNO but without DREADD expression) were not different from unmanipulated controls, indicating an absence of major off-target effects. Lastly, one might worry that activity-suppression via DREADDs is non-physiological and thus might not activate the same homeostatic mechanisms that are induced by sensory manipulations. This concern is mitigated by the observation that DREADD-induced synaptic and intrinsic homeostatic plasticity are phenotypically similar and rely on the same molecular pathways as homeostatic changes induced by pharmacological and sensory deprivation (Lambo and Turrigiano, 2013; Gainey et al., 2015; Tatavarty et al., 2020).

Ample evidence shows that sensory deprivation can induce many forms of homeostatic plasticity in a cell-type- and layer-specific manner (Feldman, 2009; Sanes and Kotak, 2011; Turrigiano, 2011). Despite these past efforts, our understanding of the developmental regulation of homeostatic plasticity remains nascent. One major reason is that different studies have used disparate paradigms (optic nerve block, lid suture, retinal lesions, enucleation) across ages and cell types, which precludes direct comparison of results from different studies. For example, synaptic scaling induced by optic nerve block is absent in L2/3 until CP onset (Desai et al., 2002), but whether it persists into adulthood has been difficult to test, mainly because visual deprivation has fundamentally different effects on V1 activity after the CP ends (Sawtell et al., 2003; Sato and Stryker, 2008). Interestingly, monocular enucleation in adult mice induces synaptic scaling in L5, but not in L2/3, pyramidal neurons (Barnes et al., 2015). In contrast, we found robust synaptic scaling in adult L2/3 pyramidal neurons following DREADD-mediated activity suppression. This demonstrates that the ability to express synaptic scaling persists in these neurons into adulthood, and suggests that enucleation either modulates the activity of L2/3 pyramidal neurons insufficiently to induce scaling, or possibly induces additional forms of plasticity that mask the homeostatic response.

Lid suture induces synaptic scaling and intrinsic homeostatic plasticity in tandem in L2/3 pyramidal neurons during the CP (Maffei and Turrigiano, 2008b; Lambo and Turrigiano, 2013), but neither L2/3 nor L5 adult pyramidal neurons express intrinsic homeostatic plasticity following bilateral retina lesion or monocular enucleation (Keck et al., 2013; Barnes et al., 2015). Here we could directly assess whether L2/3 pyramidal neurons lose the ability to express intrinsic homeostatic plasticity after the CP ends. While DREADD-mediated activity suppression induced robust intrinsic plasticity during the CP, it was completely absent in adult neurons, even after an extended period of activity suppression. Our data suggest that, unlike synaptic scaling, intrinsic homeostatic plasticity in L2/3 is tightly coupled to the CP. This critical window for intrinsic plasticity has also been reported in the auditory cortex (Rao et al., 2010). Therefore, intrinsic homeostatic plasticity and synaptic scaling are subject to starkly different developmental forces.

Like excitatory synapses, inhibitory synapses are also subject to homeostatic regulation. In the adult mouse V1, pyramidal neurons in both L2/3 and L5 show reduced mIPSC frequency following visual deprivation (Keck et al., 2013; Barnes et al., 2015; Gao et al., 2017), suggesting that excitatory neurons receive less overall inhibitory input. Intriguingly, we also observed a substantial decrease in the mean mIPSC frequency in adult L2/3 pyramidal neurons following DREADD-mediated activity suppression. This reduction in frequency could reflect inhibitory synapse loss, as reported in previous studies (Chen et al., 2011; Keck et al., 2011; van Versendaal et al., 2012). During the CP, visual deprivation-induced disinhibition in V1 is transient, and precedes and possibly creates a permissive environment for subsequent excitatory plasticity (Kuhlman et al., 2013, Hengen et al., 2013). In contrast, here in adult V1 we found that changes in excitation and inhibition occurred in parallel. The fact that this disinhibition is still present when excitatory scaling up has already been elicited suggests that it serves a purpose other than facilitating excitatory plasticity. Finally, we found that the homeostatic regulation of inhibition was also developmentally regulated. It was only observed in adults, when intrinsic plasticity was missing, but not in juveniles, when excitatory synaptic scaling and intrinsic plasticity were both present. These results support the idea that individual homeostatic mechanisms are modular, and can be turned on and off in the same cell type at different developmental stages.

Neural circuits are endowed with a diverse repertoire of homeostatic plasticity mechanisms to maintain stability during learning and development (Turrigiano, 2011). A fundamental challenge is to resolve how these mechanisms are regulated across different developmental stages to achieve the appropriate homeostatic outcome. Past efforts to answer this question have been hindered by the lack of a consistent activity manipulation method that can induce comparable changes in the same type of neurons at different ages. Here, we report a DREADD-mediated approach to bypass this limitation. Our data show that L2/3 pyramidal neurons engage strikingly distinct sets of homeostatic mechanisms in juvenile and adult mice. These mechanisms exhibit unique developmental profiles, supporting the hypothesis that individual homeostatic mechanisms subserve specific functions and will be recruited appropriately to meet changing developmental demands.

## Acknowledgments

This work was supported by NIH grant R35 NS111562 (G.G.T.). We thank Zhe Meng for technical support and all members of Turrigiano laboratory for helpful discussions.

